# α-hemolysin of *Staphylococcus aureus* impairs thrombus formation

**DOI:** 10.1101/2021.11.11.468205

**Authors:** Kristin Jahn, Stefan Handtke, Raghavendra Palankar, Thomas P. Kohler, Jan Wesche, Martina Wolff, Janina Bayer, Christiane Wolz, Andreas Greinacher, Sven Hammerschmidt

## Abstract

Toxins are key virulence determinants of pathogens and can impair the function of host immune cells including platelets. Insights into pathogen toxin interference with platelets will be pivotal to improve treatment of patients with bacterial bloodstream infections. In this study, we deciphered the effects of *Staphylococcus aureus* toxins α-hemolysin, LukAB, LukDE and LukSF on human platelets and compared the effects with the pore forming toxin pneumolysin of *Streptococcus pneumoniae*. In contrast to pneumolysin, α-hemolysin initially activates platelets as indicated by CD62P and αIIbβ3 integrin expression, but the resulting pores also induce alterations in the phenotype of platelets and induce apoptosis of platelets. The presence of small amounts of α-hemolysin (0.2 µg/mL) in whole blood abrogates thrombus formation indicating that in systemic infections with *S. aureus* the stability of formed thrombi is impaired. This might be of high clinical relevance for *S. aureus* induced endocarditis of the aortic valves. Stabilizing the thrombi by inhibiting α-hemolysin induced impairment of platelets likely reduces the risk for septic (micro-)embolization. However, in contrast to pneumolysin, α-hemolysin induced platelets damage could not be neutralized by intravenous immune globulins. In contrast to α-hemolysin, *S. aureus* bi-component pore forming leukocidins LukAB, LukED and LukSF do not bind to platelets and had no significant effect on platelet activation and viability.

Main point 1: α-hemolysin forms pores in platelets, which first activate but then result in apoptosis and impairs thrombus formation and stability

Main point 2: Polyvalent immunoglobulins do not neutralize the mode of action of the toxin

## Introduction

Platelets play an important role in haemostasis and vessel repair. They represent the smallest immune cells in humans and express *e*.*g*., Toll-like and complement receptors on their surface, thereby recognising bacterial pathogens via pathogen associated molecular patterns. Sensing and interaction of bacteria leads to platelet activation and release of antimicrobial peptides.^1^ Platelet activation can either be direct via secreted proteins or surface associated bacterial proteins or indirect via bridging molecules of the extracellular matrix (ECM).^2-4^

*Staphylococcus aureus* (*S. aureus*) and *Streptococcus pneumoniae* (*S. pneumoniae*; pneumococci) are Gram-positive facultative pathogens colonizing often asymptomatic the human upper respiratory tract. *S. aureus* is able to disseminate from the nasopharynx in other host compartments and can cause severe invasive diseases like pneumonia, infective endocarditis and bacteraemia, which can lead to organ damage and sepsis.^5,6^ Similar, pneumococci can overcome the host epithelial barrier and invade deeper host compartments and enter the blood. This causes invasive diseases like pneumonia, septicaemia, or meningitis. During dissemination via the bloodstream, bacteria get in close contact with circulating platelets. We and others have previously demonstrated the ability of *S. aureus* to activate platelets either directly via surface associated or secreted proteins (Eap, FLIPr, CHIPS, AtlA-1, α-hemolysin) or indirectly, involving host extracellular matrix (ECM) proteins.^3,7^ Pneumococci were shown to at least indirectly activate platelets via ECM proteins.^8,9^ Recently, we have shown that pneumococcal pneumolysin, a cholesterol-dependent cytolysin, does not activate but kills platelets by oligomerization on the cell and formation of pores.^10^ This may contribute to progression of pneumonia to acute respiratory distress syndrome (ARDS).^10^ α-hemolysin (Hla), released by *S. aureus* is also a pore-forming toxin. Besides its role in disrupting epithelial barriers, Hla has been described to directly activate human platelets, leading to platelet aggregation.^3,11^ Hla binds to the metalloprotease ADAM10, which is expressed on platelets.^12,13^ In contrast to pneumolysin pores (diameter of 40-50 nm), pores formed by Hla are significantly smaller (diameter of 1-4 nm).^14^ Besides Hla, *S. aureus* expresses further pore-forming toxins, the bi-component pore forming leukocidins LukSF, also referred to as Panton-Valentine Leukocidine (PVL), LukED, and LukAB (also known as LukGH).^15^ These leukocidins multimerize after binding to the membrane of the respective target cell, which results in pore formation and finally host cell death. Neutrophils and other cells of the innate immune response have been shown to be the main targets of the Luk toxins. ^16-19^ So far, only indirect effects of leukocidins on platelets have been described, which result from the destruction of neutrophils and other leukocytes.^20^ In this study, we investigated the effects of staphylococcal Hla and pore forming leukocidins on platelet activation, aggregation, viability and clot stability and compared the results with effects caused by pneumococcal pneumolysin. Gaining further insight into how bacterial toxins interfere with platelet functions is essential to improve treatment of patients suffering from systemic bacterial infections.

## Methods

### Ethics

The use of whole blood and washed platelets from healthy adult individuals was approved by the Ethics Committee of the University Medicine Greifswald (BB 044/18). All volunteers gave written informed consent in accordance with the Declaration of Helsinki. All experiments were carried out in accordance with the approved guidelines.

### Bacterial toxins

We used pneumococcal pneumolysin (Ply, 53 kDa) and *S. aureus* α-hemolysin (Hla, 33 kDa) (kindly provided by Jan-Peter Hildebrandt, University of Greifswald) recombinantly produced as described recently.^10,21^ The components LukS (33 kDa) and Luk F (34 kDa) of the pore-forming bi-component Panton-Valentine Leukocidin PVL were heterologous expressed in *E. coli* BL21 pCG 94 LukS and *E. coli* BL21 pCG142 LukF, respectively. To purify LukS and LukF Protino® Ni-TED 2000 columns were loaded with the *E. coli* cell lysate, washed 3 times with 20 mM imidazole buffer and proteins were eluted with 500 mM imidazole buffer. After verification of purity by SDS-PAGE followed by Coomassie brilliant blue R-250 staining, the proteins were dialyzed against PBS. Luk A and Luk B were heterologously expressed and purified as described elsewhere.^22^ Leukocidins E (ab190128) and D (ab190423) were purchased from Abcam (Berlin, Germany).

### Antibodies, reagents

We used the following antibodies: neutralizing mouse monoclonal anti-α-hemolysin IgG [8B7] (ab190467; Abcam, Cambridge, USA; using a rabbit red blood cell lysis assay EC^50^ of ab190467 for neutralization of 0.3 μg/mL of α-hemolysin was determined to be 0.676 μg/mL), PE-Cy5-labelled monoclonal mouse anti-human CD62P, FITC-labelled mouse PAC-1 antibodies recognizing activated α^IIb^β^III^ (CD41/CD61) (BD Bioscience, Franklin Lakes, USA), RealTime-Glo™ MT Cell Viability Assay (Promega, Madison, USA), FITC-labelled mouse anti-human CD42a (BD Biosciences, Franklin Lakes, USA), Alexa Fluor 647-labelled monoclonal mouse anti-human CD62P (P-Selectin) antibody (Clone AK4, BioLegend, San Diego, CA, USA), Alexa Fluor 647-labelled goat anti-mouse IgG (GAMIG AF-647) (Abcam, Cambridge, MA, USA) and human polyvalent immunoglobulin preparations (IVIG; IgG-enriched Privigen; CSL Behring, Marburg, Germany). Mouse polyclonal anti-LukS and anti-LukF antibodies were generated by routine immunization of mice with heterologously expressed LukS or LukF. Female CD-1 mice (Charles River Laboratories, Sulzfeld, Germany) were immunized intraperitoneally with 100 µl of a 1:1 emulsion containing 50 µg recombinant protein LukS or LukF and incomplete Freund’s adjuvant (Sigma-Aldrich, Taufkirchen, Germany). Mice were boosted with an emulsion of protein and incomplete Freund’s adjuvant at day 14 and 28 and bled after six weeks. Specificity of polyclonal antibodies was verified by immunoblot analysis (data not shown). We also used the following reagents: FAM-FLICA caspase 3/7 assay kit from ImmunoChemistry (Hamburg, Germany), Thrombin (Sigma Aldrich, Darmstadt, Germany), Convulxin (Enzo Life Sciences, Lausen, Switzerland), Ionophore – (Sigma Aldrich, Darmstadt, Germany), von Willebrand factor (vWF) (Merck, Darmstadt, Germany), Ristocetin (Mölab, Langenfeld, Germany), Annexin V (BioLegend, Koblenz, Germany), recombinant anti-Bcl-2 antibodies (AF647, Abcam, Berlin, Germany), Triton -X-100 (Sigma-Aldrich, St. Louis, USA).

### Flow cytometry-based platelet activation assay, toxin treatment of platelets and toxin neutralization

We performed all activation assays with washed platelets in Tyrode’s buffer containing Ca^2+^ and Mg^2+^ with PBS using CD62P expression as activation marker as described.^10^ In platelet activation assays with toxins, we treated platelets for 4 min with 300 ng/mL of pneumolysin or 0.02 µg/mL, 0.2 µg/mL, 2.0 µg/mL, or 20 µg/mL of α-hemolysin, LukAB, LukED or LukSF (for each pair equimolar amounts of the single leukocidins were used) followed by 5 min treatment with 20 µM TRAP-6. In neutralization experiments, we preincubated pneumolysin or α-hemolysin for 20 min at RT with 1 mg/mL human intravenous immunoglobulin (IVIG: pharmaceutical human IgG; Privigen®; CSL Behring, Marburg, Germany), or increasing concentrations of a mouse monoclonal [8B7] antibody against α-hemolysin (ab 190467; Abcam).

We measured CD62P expression using a FACSCalibur™ (Becton Dickinson) flow cytometer and CellQuestPro 6.0. We then predefined by forward-sideward-scatter a platelet gate based on measurements with CD61 positive platelets and analysed in the gated region 20,000 events for fluorescence. The value for platelet activation was calculated as the geometric mean fluorescence intensity (GMFI) of the gated population multiplied by the percentage of CD62P-positive labelled platelets.^10^

### Flow cytometry-based analysis of protein binding to human platelets

We incubated washed human platelets with human BD Fc Block™ (BD Biosciences), to prevent unspecific binding to platelet FcγRIIa; added increasing concentrations of pneumolysin, α-hemolysin, or LukSF for 10 min at 37°C, followed by fixation with PFA/PBS (pH 7.4) at a final concentration of 2% at RT for 20 min. Binding of toxins to platelets was measured using antibodies against pneumolysin (Streptavidin-Alexa Fluor 488, Dianova, Hamburg, Germany), α-hemolysin, PVL (1h; RT), and with Alexa Fluor 488 conjugated secondary antibodies for Hla and PVL (30 min; RT); using a FACSCalibur™ (Becton Dickinson) flow cytometer and CellQuestPro 6.0.

### Platelet preparation, light transmission aggregometry, live/dead staining, release of intracellular calcium, immunofluorescence staining, thrombus formation assay and western blotting

We performed platelet preparation, light transmission aggregometry, LIVE/DEAD staining, detection of Ca^2+^ released from internal stores, immunofluorescence staining, *ex vivo* thrombus formation in whole blood under shear, and western blotting as described.^10,23,24^ Details are provided in the Supplement.

### Determination of apoptosis markers

We determined platelet caspase activity, expression of Bcl-2, and exposure of phosphatidylserine (PS) as apoptosis markers. Washed human platelets were incubated in a volume of 25 µl with thrombin (10 U/mL), TRAP-6 (20 µM) and convulxin (100 ng/mL), Ionophore (10 µM) or von Willebrand factor (vWF; 20 µg/mL) and Ristocetin (1.5 mg/mL) as controls as well as with increasing concentrations of pneumolysin (3.0 – 300 ng/mL), Hla (0.2 – 20 µg/mL), or PVL (0.2 – 20 µg/mL).

We determined caspase activity using the FAM-FLICA caspase 3/7 assay kit from ImmunoChemistry (Hamburg, Germany) according to manufacturer’s instructions. In brief, 0.8 µl FLICA solution was added to the samples after toxin incubation and samples were then incubated for 45 min at 37°C in the dark. Afterwards, we added 100 µl apoptosis wash buffer, incubated samples for 7 min, centrifuged (650 x g, 7 min, RT) and measured them by flow cytometry (Cytomics FC500, Beckman Coulter, USA) after resuspension in Tyrode’s buffer. To determine Bcl-2 expression, all samples were fixed with 0.5% PFA for 20 min at RT and then centrifuged (650 x g, 7 min, RT). Platelets were then permeabilized with 0.25% saponin for 30 min and stained using recombinant anti-Bcl-2 antibodies (AF647, Abcam, Berlin, Germany) for 30 min before being measured by flow cytometry. PS-exposure was determined by Annexin V binding. We stained platelets with 5 µL Annexin V (BioLegend, Koblenz, Germany) in Annexin V binding buffer (BioLegend) containing 50 U/mL hirudin for 20 min (RT, in the dark) and measured them by flow cytometry.

### Statistics

We performed statistical analysis using GraphPad Prism (version 5.01), unless otherwise indicated. We show the data as scatter plots and include median, minimal and maximal values including median and interquartile range. We analysed the data using the nonparametric Friedman test followed by a Dunn’s multiple comparison post-test. We considered a *p*-value <0.05 to be statistically significant.

## Results

### Pneumolysin and α-hemolysin but not PVL bind to human platelets

Binding assays conducted with different amounts showed that Hla and pneumolysin bound dose-dependently to platelets in the range of 0.02 – 20 µg/ml (Hla) or 0.003 – 3.0 µg/mL (pneumolysin), respectively (Figure 1A,B), while PVL (LukSF) did not (Figure 1C).

**Figure 1.**
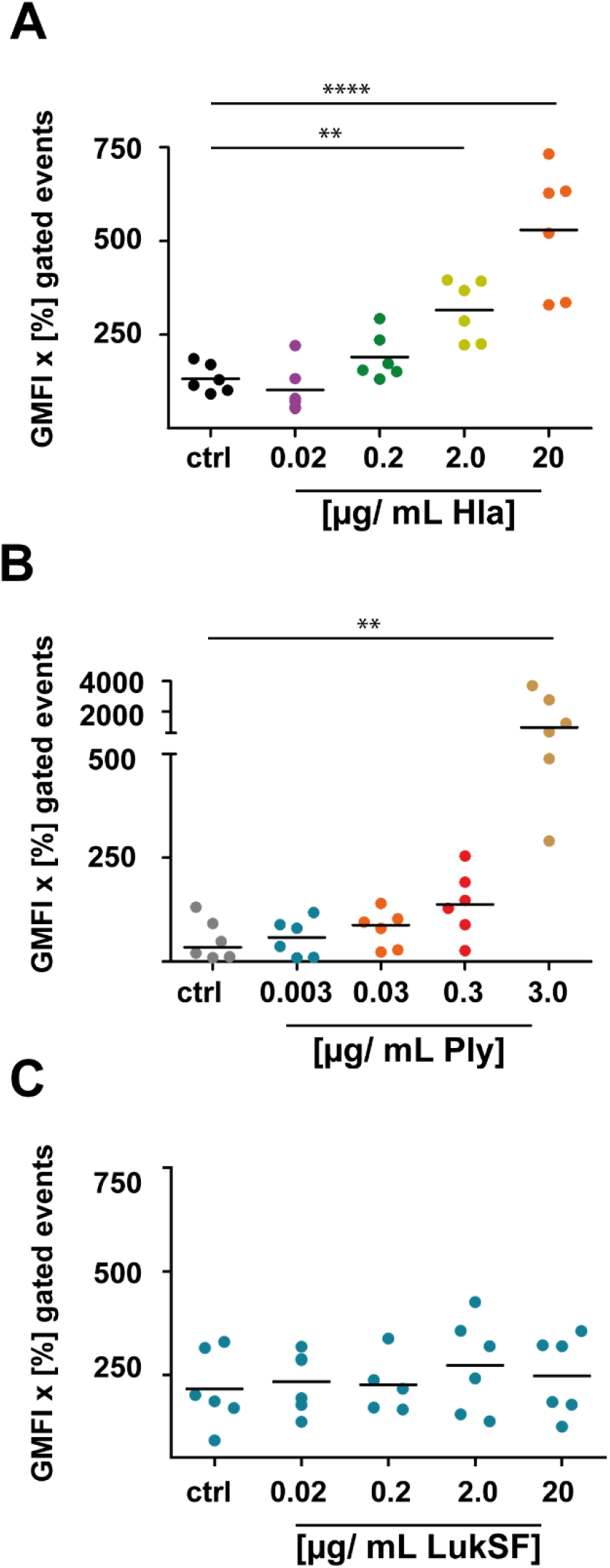
Pneumolysin and α-hemolysin bind directly to human platelets. Washed platelets of a defined set of healthy human donors were incubated with increasing concentrations of pneumolysin (Ply 0.3 to 300 ng/mL), α-hemolysin (Hla 0.02 to 20 µg/mL) and LukSF ((PVL) 0.02 to 20 µg/mL), fixed and stained with antibodies directed against the toxins (Hla, LukSF) or their Strep-tag (Ply). To exclude binding of the antibodies to the platelet Fcγ receptor, the receptor was blocked with Human BD Fc Block™. Binding events were detected by flow cytometry. The data are presented as geometric mean of fluorescence intensity (GMFI) of the positive gated events multiplied with the percentage of positive gated events in the dot plots. (A) Platelets were treated with PBS (grey) or increasing concentrations of Hla for 10 min. PBS treated platelets were used as a negative control. The staphylococcal Hla binds dose-dependently to washed human platelets, starting at a concentration of 0.2 µg/ml. (B) Platelets were treated with PBS (grey) or increasing concentrations of pneumolysin for 10 min. PBS (grey) treated platelets were used as a negative control. Binding of pneumococcal pneumolysin to human platelets was detectable starting at a concentration of 30 ng/mL. (C) Platelets were treated with PBS (grey) or increasing concentrations of LukSF (PVL) for 10 min. PBS (grey) treated platelets were used as a negative control.

### α-hemolysin but not bi-component leukocidins activate platelets

To investigate platelet activation by bacterial toxins, we treated washed human platelets with the toxins. After 10 min of incubation, Hla ≥ 2.0 µg/ml and Ply ≥ 30 ng/ml increased the CD62P signal of washed platelets. TRAP-6 stimulation did not further increase this CD62P signal. In contrast, PVL, LukAB, and LukED had no effects on platelet CD62P expression nor on platelet responsiveness to TRAP-6 stimulation (Figure 2A).

**Figure 2.**
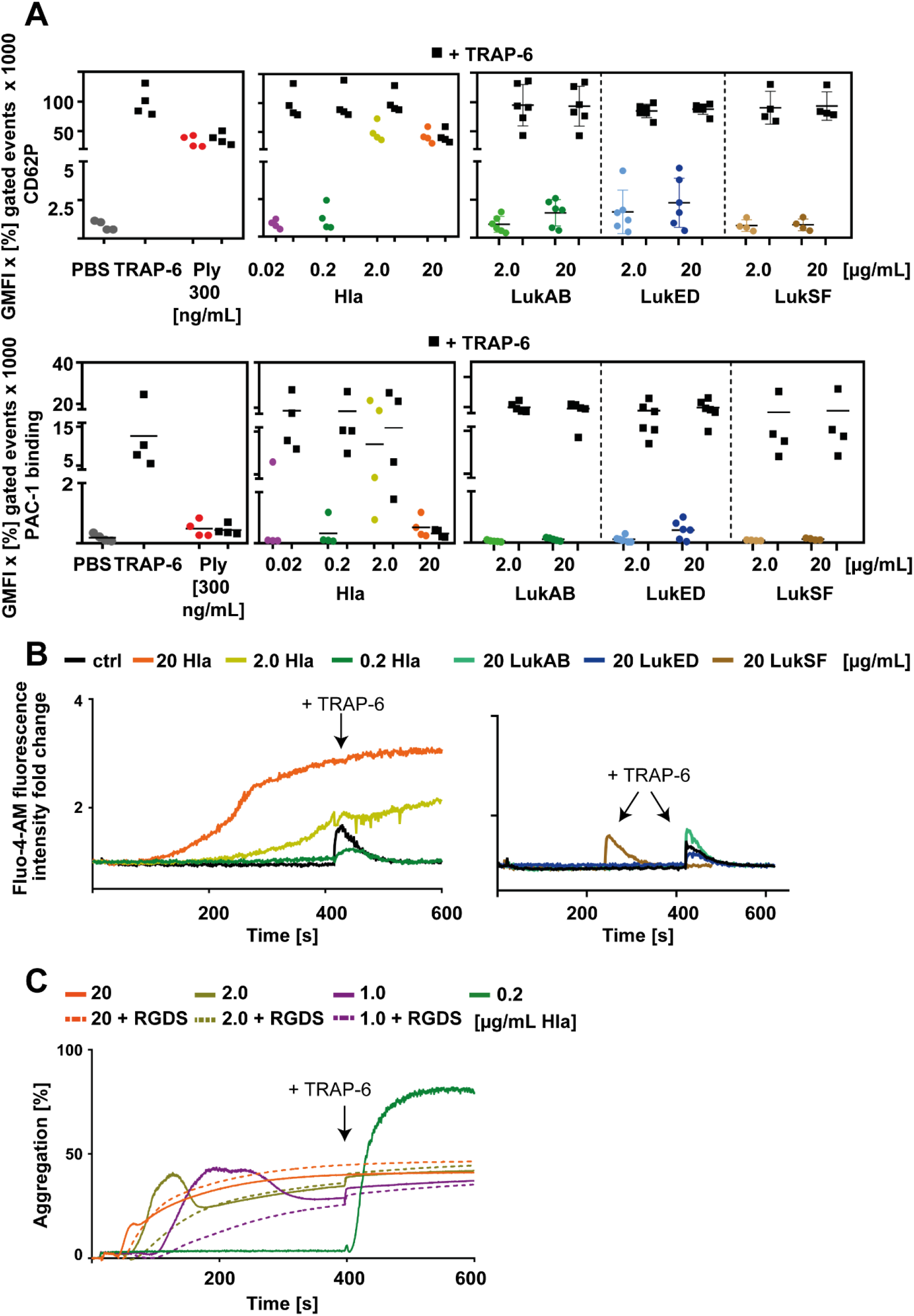
α-hemolysin and pneumolysin interfere with platelet function but with different mechanisms. Washed platelets of a defined set of healthy human donors were incubated with increasing concentrations of pneumolysin (Ply 0.3 to 300 ng/mL), α-hemolysin (Hla 0.02 to 20 g/mL) LukAB, LukED and LukSF (PVL) for 10 min. (A) CD62P (upper panel) and PAC-1 binding (lower panel) were used as activation markers and detected by flow cytometry, using a PE-Cy5 labelled P-selectin antibody and a FITC labelled anti-human GPIIbIIIa antibody (PAC-1). PBS was used as negative control and 20 µM TRAP-6 as positive control. Platelets were incubated with the toxins for 10 min. Alternatively, after 5 min of incubation with the toxins, the platelets were additionally stimulated with 20 µM TRAP-6 for 5 min to proof functionality. The data are presented as geometric mean of fluorescence intensity (GMFI) of positive gated events multiplied with the percentage of positive gated events in the dot plots. (B) Prior to treatment with pneumolysin, Hla, LukA/B, LukD/E or LukSF (PVL), intracellular Ca^2+^ of washed platelets was labelled with Fluo-4-AM for 30 min. After incubation with increasing concentrations of the indicated toxins the kinetics of Ca^2+^ release was measured and values are given as fold change compared to NaCl control. (C) Platelet aggregation was measured using light transmission aggregometry. Hla concentrations ≥ 2 µg/ml induced an increase in light transmission, but platelets where no longer responsive to 20 µM TRAP-6, which was subsequently added after 6 min of incubation.

Platelets incubated with > 0.2 µg/mL Hla showed an increased signal for αIIbβ3 integrin activation (Figure 2A), while platelets incubated with 0.3 µg/mL Ply did not. At concentrations > 0.2 µg/mL Hla induced release of intracellular calcium (Figure 2B) and increased light transmission in the aggregometer (Figure 2C). While the curves for Ca^2+^ release gradually increased (Figure 2B), a partly reversibly change in light transmission was observed in the aggregometer. We therefore measured the change in light transmission in the presence of RGDS, which inhibits platelet aggregation. Any change in light transmission measured in the presence of RGDS is caused by platelet lysis. Overlay of the curves reveals the following sequence of events (Figure 2C). Hla first induces platelet activation and aggregation (first peak of the curve) in parallel to calcium influx. Then platelets are killed by the toxin, start to disaggregate and lysis occurs. The aggregation curve overlays the curve of platelet lysis induced change of light transmission (measured in the presence of RGDS) after about 180 sec for 2.0 µg/mL Hla and after about 400 sec for 1.0 µg/mL Hla. LukSF, LukAB or LukED did not induce calcium release or an increase in light transmission in the aggregometer (Figure 2B and data not shown).

### Platelets are killed by prolonged exposure to α-hemolysin

Previously, we demonstrated that pneumolysin does not cause platelet activation but directly kills platelets by formation of large pores (40-50 nm). The CD62P signal induced by pneumolysin results from antibody diffusion into the cytoplasm through the pores and intracellular CD62P staining instead of platelet activation.^10^ From the experiments described above we concluded that the initial increase in CD62P and the first peak of an increase in light transmission in aggregometry of Hla treated platelets represents platelet activation. However, like pneumolysin, Hla also forms pores in cell membranes, but the pore size is much smaller (1.5-2.0 nm) and theoretically too small for antibodies to pass through. By CLSM we visualized CD62P on the surface of platelets in response to Hla (Figure 3A). Triton X-100 (control for intracellular CD62P staining) treated platelets were permeabilized and intracellular CD62P was stained. TRAP-6 (control for platelet membrane CD62P staining) incubated platelets showed, similar to Hla treated platelets, CD62P on the surface (Figure 3A). However, Hla treated platelets were enlarged and swollen compared to the TRAP-6 control, suggesting that Hla induces loss of platelet membrane integrity and subsequently loss of platelet function.

**Figure 3.**
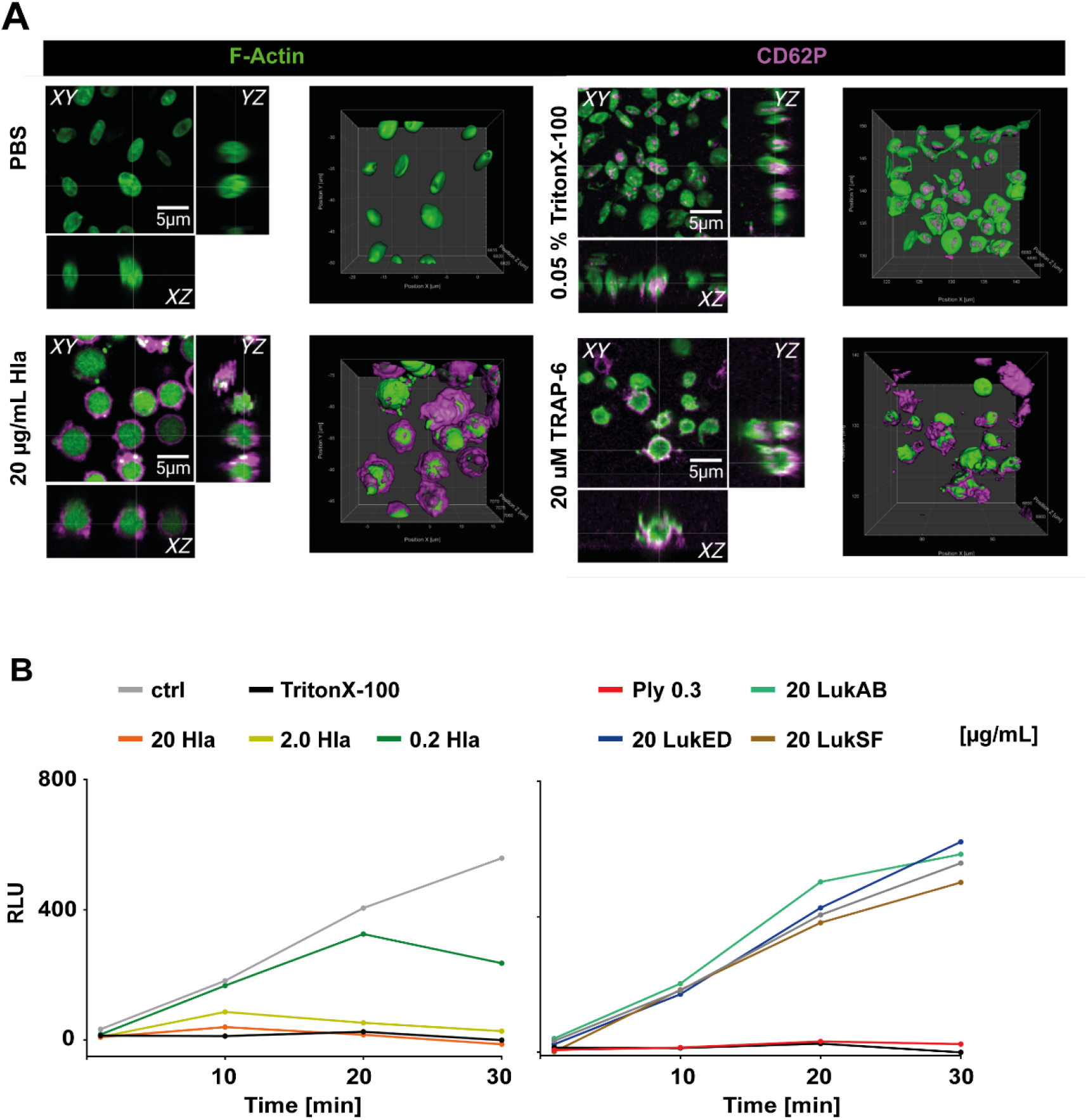
Treatment of human platelets with α-hemolysin leads to staining of surface associated CD62P. (A) α-hemolysin (Hla) treated platelets were stained for F-actin (green) and CD62P (magenta). Platelets were not permeabilized. Orthogonal views of confocal Z-stacks and 3D iso-surface rendering of platelets are shown. Platelets treated with 20 µg/mL Hla show distinct extracellular staining of CD62P comparable with only TRAP-6 treated platelets. TRAP-6 was used as control for surface associated CD62P and TritonX-100 as control for intracellular CD62P staining. (B) Kinetics of platelet viability measured with the RealTime-Glo™ MT Cell Viability Assay (Promega). PBS was used as viability control and Triton X-100 to induce platelet death. Increasing concentrations of Hla, LukA/B, LukD/E, LukSF (PVL) and 300 ng/mL pneumolysin were incubated for 30 min with washed platelets. One minute after mixing of platelets and toxins the measurement started.

We also concluded from the platelet aggregometry experiment that after initial activation, platelets are killed. We therefore measured the viability of platelets exposed to different concentrations of toxins over 30 min. Pneumolysin was used as “killing” control. Low concentrations of Hla (0.2 µg/mL) reduced platelet viability after 20 min. In contrast, higher Hla concentrations killed platelets rapidly (Figure 3B). Only at 0.02 µg/mL Hla platelet viability remained unaffected and RLU values increased similar to the PBS control (data not shown).

### α-hemolysin and pneumolysin induce apoptosis in human platelets

As pneumolysin^10^ and Hla differ in their initial effects on platelets, we asked whether these toxins differ in their capability and mechanism to induce cell death. Hla and pneumolysin strongly induced phosphatidylserine (PS)-exposure of platelets. This signal was comparable to or even higher than the signal obtained for the positive controls ionophore and convulxin (Figure 4A). Both toxins, pneumolysin and Hla dose-dependently increased caspase-3/7 activity (Figure 4B) in platelets, but did not increase Bcl-2 expression (Figure 4C). This suggests that both toxins induce apoptosis by activating effector caspases (Figure 4B). The toxin concentrations showing activation of cell death and apoptosis markers correspond to the concentrations inducing a loss of platelet function (Figure 2 and 3).

**Figure 4.**
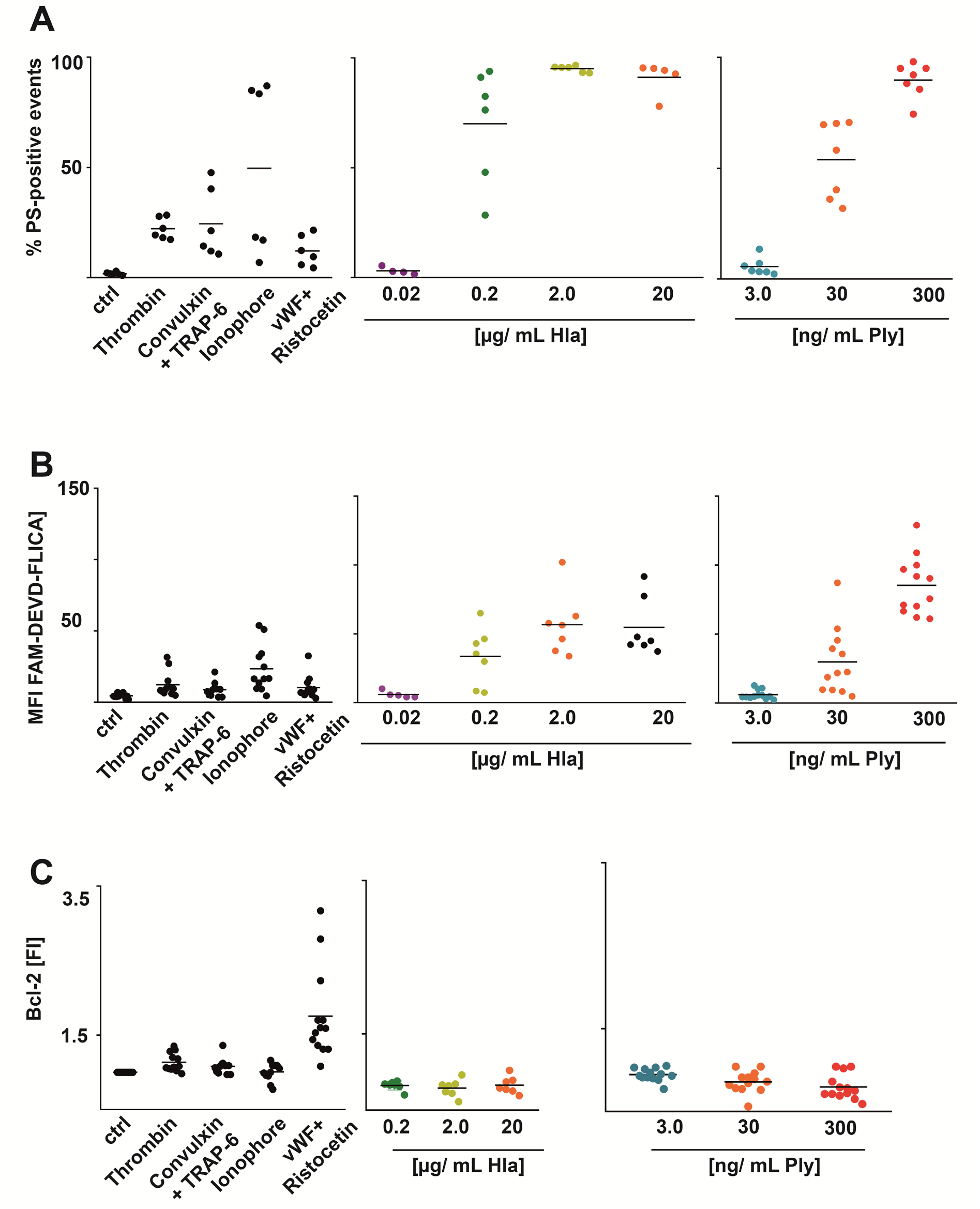
Platelets treated with α-hemolysin and pneumolysin are positive for apoptosis markers. Washed human platelets were incubated with increasing concentrations of α-hemolysin (Hla) and pneumolysin (Ply). The analysis of the apoptosis markers caspase activity, Bcl-2 expression and phosphatidylserine (PS) exposure was measured using flow cytometry. For all experiments thrombin, convulxin/Trap-6, Ionophore and vWF (von Willbrand factor)/Ristocetin were used as positive controls. PBS was used as negative control. (A) PS-exposure was determined by Annexin V binding. Values are given as percent of positive events. Treatment with Ply and Hla leads to PS-exposure in a concentration dependent manner. (B) Caspase activity was measured by fluorescent labelling of active caspase 3 and 7 in Ply- or Hla treated human platelets. Values are given as mean fluorescent intensities and show a dose dependent increase after pneumolysin- or Hla treatment. (C) Bcl-2 expression was determined using a recombinant anti-Bcl-2 antibody. After treatment with Hla or Ply, platelets were fixed and analyzed for Bcl-2 expression using flow cytometry. Values are given as fluorescence intensities.

### Polyvalent immunoglobulin preparations did not inhibit platelet damage by α-hemolysin

Recently, we showed that IVIG or specific anti-pneumolysin antibodies prevent killing of platelets by pneumolysin.^10^ Based on these findings, we assumed that IVIG and a specific neutralizing monoclonal IgG antibody targeting Hla also have the potential to inhibit loss of platelet function and cell death. Both, IVIG and a mouse anti-α-hemolysin antibody recognise purified Hla (Figure S2). However, they did neither prevent CD62P expression nor loss of platelet viability in response to Hla (Figure 5A, B; Figure S2).

**Figure 5.**
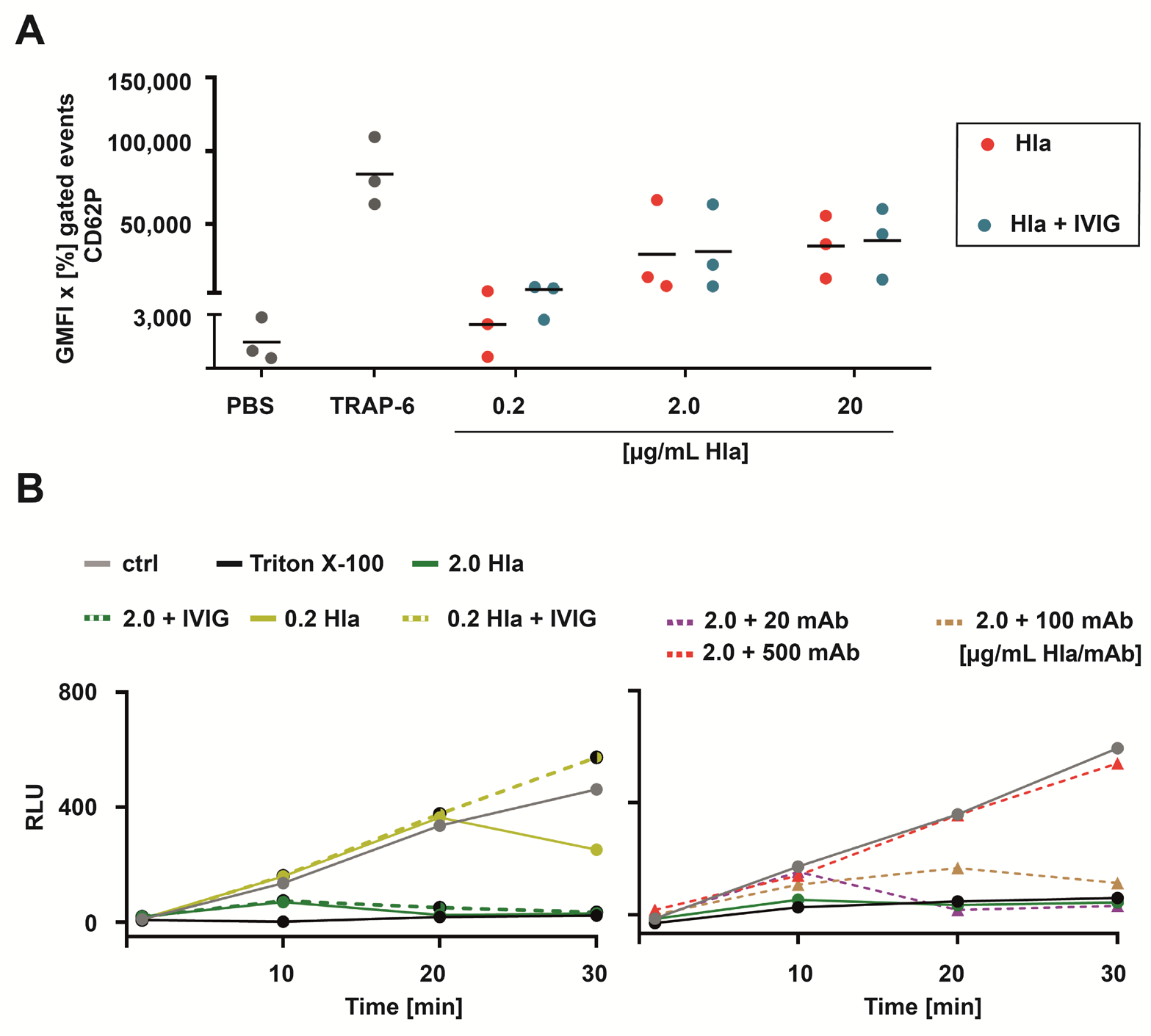
IVIG does not neutralize α-hemolysin. α-hemolysin (Hla) was treated with IVIG (1mg/ml) or a specific mAb for 20 min before incubation with washed human platelets of healthy human donors. (A) Polyvalent human immunoglobulins (IVIG; 1 mg/ml; human IgG, Privigen) did not neutralize the increased CD62P signal after Hla treatment. The data are presented as geometric mean of fluorescence intensity (GMFI) of the positive gated events multiplied with the percentage of positive gated events in the dot plots. (B) Platelet viability was only barely improved by IVIG (1 mg/ml) or a specific monoclonal anti-Hla antibody. After treatment with 0.2 µg/mL α-hemolysin, IVIG rescued the decreasing platelet viability over time, and 500 µg/mL of the anti-Hla antibodies rescued platelet viability.

### Thrombus formation under shear is abrogated by α-hemolysin

To assess whether Hla impacts the capability of thrombus formation under shear, we next perfused whole blood in the absence or presence of Hla at different concentrations. Hla at the lowest concentration of 0.2 µg/mL significantly reduced thrombus formation and area covered by thrombi by more than 50% (*P*<0.001) compared to the control (Figure 6A). Similarly, at higher concentrations (2.0 and 20 µg/mL), Hla strongly reduced the capacity of platelets to form stable thrombi. IVIG (1 mg/mL) failed to restore the ability of platelets to form stable thrombi under shear in the presence of Hla (Figure 6B).

**Figure 6.**
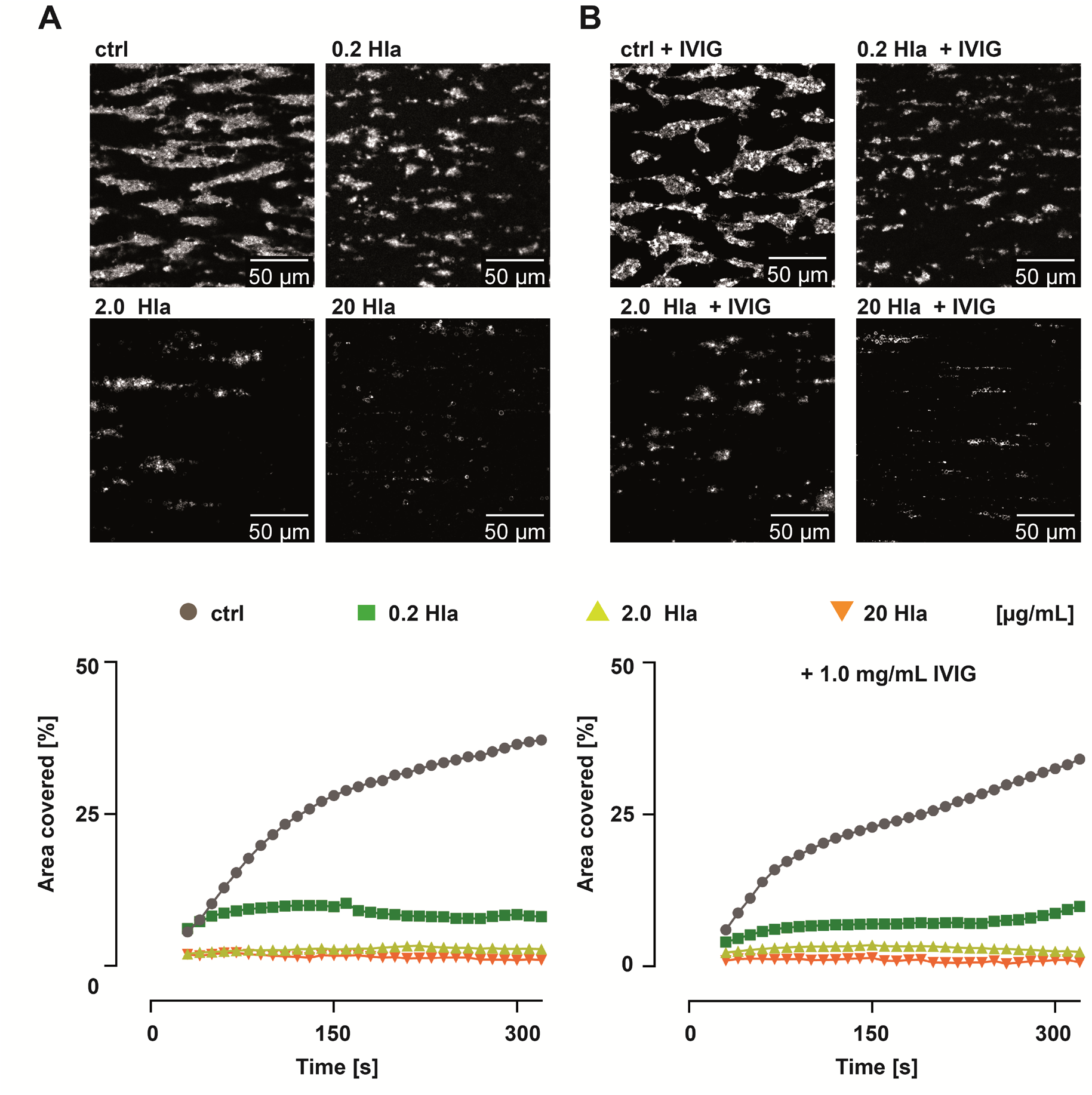
α-hemolysin induces abrogation of thrombus formation in whole blood under shear. Whole blood incubated with α-hemolysin (Hla) at 0.2, 2 and 20 µg/mL Hla was perfused over a collagen (shear 1000s^-1^) coated surface and thrombus formation was visuliazed by immunostaining of platelets with fluorescently labelled anti-CD61antibody. (A) Representative inverted gray scale images of the impact of Hla compared to the non-treated control show that Hla strongly inhibits thrombus formation under shear. (B) In the presence of IVIG (1mg/mL) thrombus formation under shear is not restored. (C) Area covered by thrombi under shear in the presence of increasing concentrations of Hla in comparison to controls and after treatment with IVIG. The data show percentage of area covered by thrombi computed from three different regions of interest from n=3 donors. Statistical analysis was performed with one-way ANOVA with Šidák multiple correction. P<0.05 was considered significant.

## Discussion

In this study we show that the *S. aureus* toxin Hla directly activates but finally kills platelets dose-dependently, while bi-component leukocidins have no direct effects on platelets.^17^ We further indicate that Hla abrogates thrombus formation in whole blood and that Hla cannot be neutralized by IVIG.

Our binding assays demonstrate that Hla binds directly to platelets, whereas the bi-component toxin LukSF (PVL) do not interact directly with platelets. LukSF is known to bind to complement receptors C5aR1 and C5aR2 on leukocytes.^17,25^ Lack of platelet stimulation via LukSF can be explained by the lack of C5a receptor (C5aR) exposed on the platelet surface. This is supported by a recent study showing a C5aR transcript in platelets but the receptor protein was not detected by proteomics.^26^ Similarly, LukED interacts with receptor CCR5, CXCR1, and CXCR2.^20,27,28^ Also the transcripts for CCR5, CXCR1, and CXCR2 were identified in platelets, whereas the protein was absent ^26^. LukAB had also no effect on platelet activation and aggregation in our study. LukAB binds to CD11b,^29^ which is not expressed by platelets. However, supernatants of neutrophils incubated with staphylococcal pore-forming leukocidins induce platelet activation and aggregation.^20^ Our platelet binding and activation data support the concept, that *S. aureus* bi-component leukocidins only indirectly activate platelets via leukocyte activation.^20^ The data also support that these toxins are highly receptor dependent and that the cognate receptors are not expressed on platelets obtained from healthy donors.

*S. aureus* Hla forms pores of 1.5-2.0 nm diameters and is a major virulence determinant for staphylococcal infections.^21,30,31^ Hla promotes blood coagulation via activation of human platelets. This phenomenon is independent of platelet killing,^32-34^ and consistent with the strong procoagulatory PS exposure on the platelet membrane induced by Hla. *In vivo*, intravenous injection of Hla in mice induced platelet aggregation and formation of micro-thrombi. The aggregates are retained in the liver sinusoids and kidney glomeruli, thereby causing multi-organ dysfunction.^11^

Our studies suggest that Hla acts in two steps on platelets. The small pores formed by Hla lead to calcium influx and initial platelet activation and aggregation. Evidence for platelet activation are surface-exposed CD62P and the ability of RGDS to block the initial aggregation peak. However, increasing pore formation finally kills platelets. This explains, why Hla induced platelet activation and cell death are time and concentration dependent. Higher Hla concentrations (≥ 2.0 µg/mL Hla) strongly increased CD62P expression, and killed platelets within 2 to 7 min, resulting in a total loss of platelet function. Lower concentrations (0.02 µg/mL), induced killing after 20 min of incubation. In contrast, pneumolysin forms large pores in the platelet membrane and thereby kills platelets immediately without previous activation, even at very low concentrations.^10^

We used much lower Hla concentrations (max. 0.56 µM) compared to the Hla concentrations found in patient sera (up to 60 µM).^34^ While platelet activation by various *S. aureus* proteins like Clumping factor A (ClfA), SdrE, AtlA1, CHIPS, FLIPr, and Eap including Hla is well accepted, the consequences of platelet killing by *S. aureus* has gained less attention.^3,36^ However, killing of platelets might be clinically highly relevant. One of the most feared infections of *S. aureus* is endocarditis. The biggest risk in acute endocarditis are septic thrombi causing multiple occlusions of small arteries, especially in the brain. In this regard, our finding that Hla destabilizes thrombi has major implications. Our data indicate that thrombus stabilizing treatment potentially reduces the risk of microthrombi dissemination from the infected aortic valve in *S. aureus* induced endocarditis.

We therefore addressed the question, whether platelet killing by Hla can be inhibited. Most individuals have anti-Hla IgG antibodies in their plasma. We therefore tested the potential neutralizing effect of the pharmaceutic immunoglobulin preparation IVIG on Hla, which, however, did not abrogate platelet killing by Hla. Besides IVIG, anti-Hla monoclonal antibodies might be an option. Although the monoclonal antibody tested in this study had no effect on Hla induced killing of platelets, a humanized Hla neutralizing antibody (MEDI4893*)^35^ inhibited organ damage in *S. aureus* sepsis in animal models^11^ and is well tolerated in humans^36^. The antibody was not effective in preventing *S. aureus* induced pneumonia in intensive care patients^37^, but its effects on thrombus stabilization have not been assessed up to now.

The receptor for Hla on platelets is the widely expressed metalloprotease ADAM10.^13,26^ Depletion of this receptor has been shown to prevent Hla induced cellular damage and dysfunction.^38^ Furthermore, inhibition of ADAM10 was shown to attenuate vascular injury during sepsis in mice.^39,40^ However, due to incomplete mechanistic understanding of the regulation of metalloproteases, clinical trials with metalloprotease inhibitors failed up to now.^41^

Finally we addressed, how Hla causes platelet death. Hla strongly increases caspase 3/7 activity, indicating apoptotic cell death. Bcl-2 as an anti-apoptotic signal inhibiting caspase activity was not increased.^42^ Future studies should address whether other cell death mechanisms are also involved like ferroptosis, necrosis or necroptosis.

Taken together, we demonstrate that *S. aureus* Hla but not leukocidins interplay with platelets. Hla initially activates platelets followed by cell death. Platelets undergo apoptosis, which leads to thrombocytopenia and impairment of thrombus stability. Inhibiting Hla might be a relevant factor to mitigate the risk of dissemination of septic microthrombi in *S. aureus* endocarditis.

## Acknowledgement

The authors thank Peggy Stremlow (University of Greifswald) for technical support.

## Authorship

Contribution: Kristin Jahn performed binding experiments, flow cytometry and cell viability experiments, evaluated the data, prepared the figures and wrote the manuscript;

Stefan Handtke: performed calcium assays, aggregometry and evaluated the data, prepared figures and edited the manuscript;

Raghavendra Palankar: designed and performed platelet confocal microscopy, evaluated the data, prepared the figure and edited the manuscript;

Thomas P. Kohler: contributed to the flow cytometry experiments, designed experiments, and edited the manuscript;

Jan Wesche: contributed to flow cytometry experiments, platelet function studies, managed healthy donors, and edited the manuscript;

Martina Wolff: performed apoptosis experiments

Janina Bayer: purified leukocidins LuSF and LukAB

Christiane Wolz: purified leukocidins LuSF and LukAB

Andreas Greinacher: designed the project, funding of the project, supervised the project, evaluated the data, wrote and edited the manuscript.

Sven Hammerschmidt: designed the project, funding of the project, supervised the project, evaluated the data, wrote and edited the manuscript;

All authors reviewed the final version of the manuscript

### Correspondence

Sven Hammerschmidt, Department of Molecular Genetics and Infection Biology, Interfaculty Institute of Genetics and Functional Genomics, Center for Functional Genomcis of Microbes, Universität Greifswald, Felix-Hausdorff-Str 8, D-17487 Greifswald, Germany; and Andreas Greinacher, Institut für Immunologie und Transfusionsmedizin, Universitätsmedizin Greifswald, Sauerbruchstr, 17487 Greifswald, Germany.

