## Supplemental Material for "α-hemolysin of *Staphylococcus aureus* impairs thrombus formation"

**Supplemental Methods**

**Supplemental Figures S1 and S2**

### $\alpha$-hemolysin of *Staphylococcus aureus* impairs thrombus formation

Kristin Jahn[1], Stefan Handtke[2], Raghavendra Palankar[2], Thomas P. Kohler[1], Jan Wesche[2], Martina Wolff[2], Janina Bayer[3], Christiane Wolz[3,4], Andreas Greinacher[2#], Sven Hammerschmidt[1#],

[1]Department of Molecular Genetics and Infection Biology, Interfaculty Institute for Genetics and Functional Genomics, Center for Functional Genomics of Microbes, University of Greifswald, Germany

[2]Department of Transfusion Medicine, Institute of Immunology and Transfusion Medicine, University Medicine Greifswald, Germany

[3]Interfaculty Institute of Microbiology and Infection Medicine, University of Tübingen, Tübingen, Germany.

[4]Cluster of Excellence EXC 2124 "Controlling Microbes to Fight Infections", Tübingen, Germany.

### Methods

#### Platelet preparation

Platelets were purified from ACD-A anticoagulated whole blood from healthy donors, not taking any antiplatelet drugs or non-steroidal anti-inflammatory drugs (NSAIDs). We prepared platelets as described.[1] In brief, we washed platelet rich plasma (PRP) twice with Tyrode´s buffer containing 0.35% BSA, 0.1% glucose, 2.5 U/mL apyrase, 1 U/mL hirudin, pH 6.3 and resuspended the final platelet pellet in a bicarbonate based suspension buffer containing 0.35% BSA, 0.1% glucose, 0.212 M $MgCl_2$, 0.196 M $CaCl_2$, pH 7.2. Platelets were adjusted to 300,000 platelets/µL.[2]

#### Light transmission aggregometry

We resuspended washed platelets in suspension buffer and added fibrinogen to a final concentration of 2.25 mg/mL. In some experiments RGDS peptides to a final concentration of 1.16 mM, or a pneumolysin inhibiting monoclonal mouse antibody (7.5 µg/mL), polyclonal rabbit anti-pneumolysin antibodies (10 µg/mL), or a human polyvalent immunoglobulin preparations (IVIG; IgG-enriched Privigen; CSL Behring, Marburg, Germany) were added. After transfer to the aggregometer cuvette, different concentrations of pneumolysin were added to the platelet suspension after 15 s. We measured platelet aggregation as a decrease in turbidity of the medium with an APACT4 aggregometer at 500 rpm, 37°C (Haemochrom, Germany) applying APACT LPC-software. In some experiments we added 20 µM TRAP-6 after 240 s and continued measurement for further 200 s.

**Live/dead staining**

For measurement of cell viability in the presence of pneumolysin, Hla and the leukocidins LukAB, LukSF (PVL) and LukDE, we used the RealTime-Glo$^{TM}$ Cell Viability Assay kit (Promega) and measured viability for 30 min. We mixed the assay substrate 1:1 with the 2-fold concentration of pneumolysin, Hla or LukSF. In a subset of experiments, a Hla inhibiting monoclonal mouse antibody, or IVIG (Privigen; CSL Behring, Marburg, Germany) was added. Then we added the toxin-substrate mixture to washed human platelets in a 96 well plate in duplicates. After 1 min of incubation, we started shaking the plate with 300 rpm for 3 s and measured relative luminescence units (RLU) using a microtiter plate reader. We repeated shaking and measurement of luminescence every 60 s until a total measurement time of 30 min. Sample values of luminescence represent the mean of the duplicates subtracted by blank (Tyrode's buffer without platelets) of six independent experiments.

**Release of intracellular calcium**

We detected the release of $Ca^{2+}$ from internal stores to the cytoplasm by fluorescent labelling of free intracellular $Ca^{2+}$ using Fluo-4-AM (ThermoFisher, USA). We resuspended platelets in PBS without $MgCl_2$ and $CaCl_2$ (pH 7.4), adjusted them to 150,000 platelets/µL and stained them with Fluo-4-AM for 30 min in the dark at RT. After a 1:2 dilution in PBS, we carried out baseline measurements for 15 s. Afterwards, we stimulated platelets with pneumolysin (3-300 ng/mL final), hla (0.2-20 µg/mL final) or LukSF (0.2-20 µg/mL final). We measured free $Ca^{2+}$ with a Fluoroskan Ascent FL fluorometer (ThermoFisher, USA) over 7 min. In some experiments we added TRAP-6 at a final concentration of 20 µM after 250 s and the measurement was carried out for another 200 s.

**Immunofluorescence staining of platelets**

Immunofluorescence analysis of CD62P localization was done as described recently[3]. In brief, we incubated washed human platelets (300,000 cells/µL) in PBS (negative control), or 20 µM TRAP-6 (positive control), or 0.05% Triton X-100 (control for membrane disintegration) or 20 µg/mL Hla for 10 min at 37°C. After fixation with 2% PFA the samples were spun on microscopy slides using the Cytospin system (Thermo Fisher). We stained platelets with AF647 labelled CD62P antibody and 20 µM phalloidin ATTO 488. The samples were covered with 20 µL of Fluorescent Mounting Medium (ROTI Mount FluorCare HP19, Carl Roth GmbH Karlsruhe, Germany). Imaging was performed on a Leica SP5 confocal laser scanning microscope (Leica, Wetzlar, Germany) equipped with HCX PL APO lambda blue 40.0x/1.25 OIL UV objective. For detailed analysis of localization of CD62P we performed confocal Z-stacks of platelets and created orthogonal views and 3D rendering.

***Ex vivo* thrombus formation assay under shear**

Hirudinized whole blood (1 mL) was incubated with alpha-hemolysin at 0.2 µg/mL, 2 µg/mL and 20 µg/mL final concentration for 10 min.. In a subset of experiments, IVIG (Privigen; CSL Behring, Marburg, Germany) was added to hirudinized whole blood either in the absence or in the presence of alpha-hemolysin. Thrombus formation assays were performed (n=3 donors) at a wall shear rate of 1000 $s^{-1}$ on collagen-passivated surfaces (200 µg/mL HORM collagen type I from horse tendon; Nycomed) in a microfluidic parallel platelet flow chamber (on µ-Slide VI 0.1 with physical dimensions: 1 mm width, 100 µm height, and 17 mm length [Ibidi]).[3-5] Platelets in whole blood were immunofluorescently labeled with monoclonal antibody CD61-

FITC (Cat. No. 561913, RRID: AB_396094 BD Biosciences, Franklin Lakes, USA, at a final concentration of 0.125 µg/mL) to facilitate visualization of thrombus formation. Time-lapse confocal imaging was performed at intervals of 10 seconds per image on a Leica SP5 confocal laser scanning microscope (Leica) equipped with a water immersion HCX PL APO Lambda blue 40×/1.25 UV objective. FITC was exited at 488 nm with an argon laser line selected with AOTF; fluorescence emission was collected between 505 and 515 nm on a HyD. Assessment of platelet adhesion and thrombus formation to obtain the percentage area covered by thrombi ( n=3 images measuring field of view 193.75 µm2 each from different regions of interest from per donor) was performed in Fiji. All flow chamber perfusion experiments were performed according to the International Society on Thrombosis and Haemostasis Scientific and Standardization Committee (ISTH SSC) subcommittee on Biorheology recommendations.

**Western Blot**

A total of 1 µg of pneumolysin, Hla and LukSF (PVL), respectively, was separated on a 12% SDS-PAGE. The samples were blotted using a semi-dry blotting system (Trans-Blot SD Cell, Bio-Rad) on a nitrocellulose membrane. After blocking with 5% skimmed milk powder (Roth, Karlsruhe) in TBS, we used IVIG (Privigen; CSL Behring, Marburg, Germany) and a HRP-conjugated secondary antibody to detect the toxins with a ChemoCam (Intas, Science Imaging). The Immunoblot images were adjusted for brightness and contrast using Photoshop CS5 64 bit.

#### Supplemental Figures:

**Figure S1**

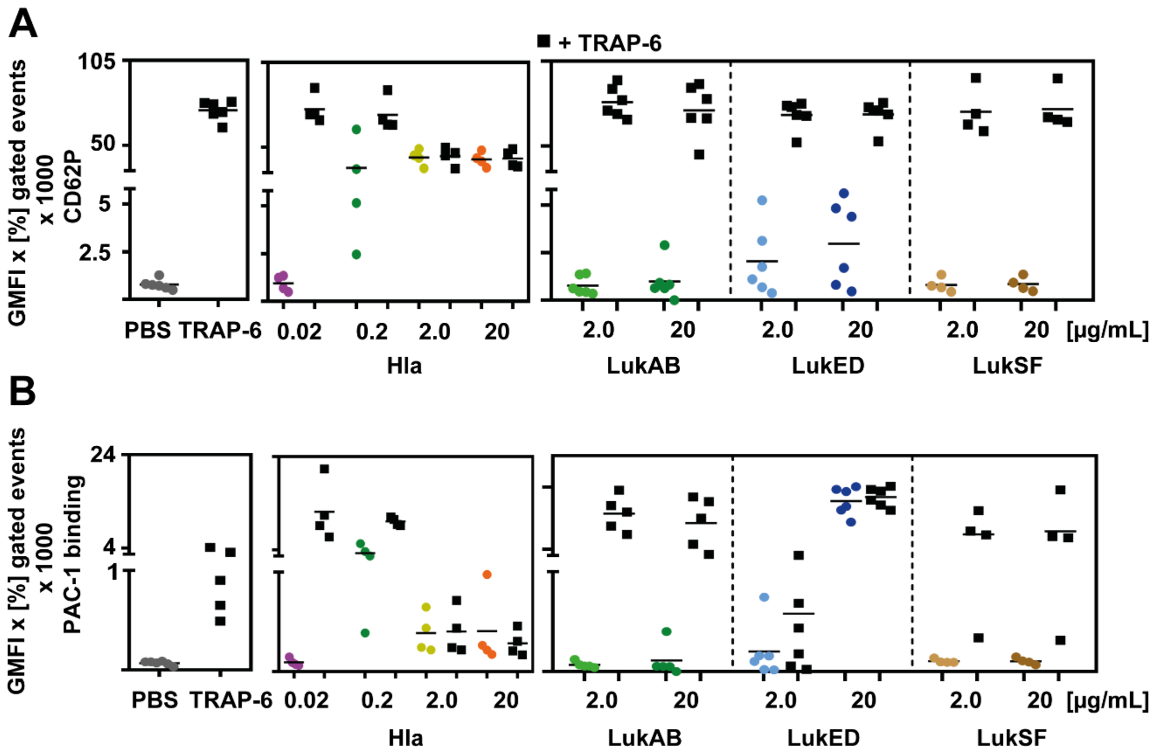


**Figure S1. Hla induced CD62P expression and PAC-1 binding over time**

Washed platelets of a defined set of healthy human donors were incubated with increasing concentrations of alpha-hemolysin (Hla, 0.02 to 20 g/mL), LukAB, LukED and LukSF (PVL) for 10 or 30 min.

(A) CD62P was used as activation marker and detected by flow cytometry, using a PE-Cy5 labelled P-selectin antibody. PBS was used as negative control and 20 µM TRAP-6 as a positive control. Platelets were incubated with the toxins for 30 min. Alternatively, after 5 min of incubation with the toxins, the platelets were additionally stimulated with 20 µM TRAP-6 for 5 min to proof functionality. The data are presented as geometric mean of fluorescence intensity (GMFI) of positive gated events multiplied with the percentage of positive gated events in the dot plots.

(B) PAC-1 binding was used as activation marker and detected by flow cytometry, using a FITC labelled anti-human PAC-1 antibody. PBS was used as negative control and 20 µM TRAP-6 as a positive control. Platelets were incubated with the toxins for 30 min. Alternatively, after 5 min of incubation with the toxins, the platelets were additionally stimulated with 20 µM TRAP-6 for 5 min to proof functionality. The data are presented as geometric mean of fluorescence intensity (GMFI) of positive gated events multiplied with the percentage of positive gated events in the dot plots. The decreased signals compared to 10 min of incubation is due to ongoing internalization of the activated GPIIb/IIIa.

#### Figure S2

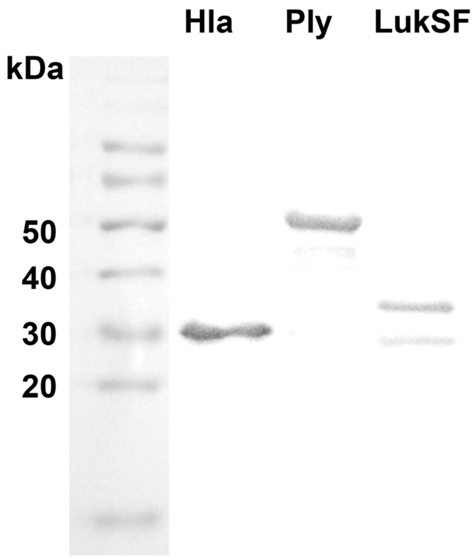


**Figure S2. IVIG recognizes pneumolysin, Hla and LukSF**

One µg of the indicated proteins or in the case of LukSF (PVL) protein mixture (equimolar ratio) were applied on a 12% SDS gel, blotted on a nitrocellulose membrane and stained with 1 mg/ml IVIG as primary antibody and a HRP coupled anti-human IgG as secondary antibody.
